# RNA Methylation dynamics regulates cancer stemness through modulation of cell migration and cell proliferation in the spheroidal model of TNBC

**DOI:** 10.1101/2025.07.22.666067

**Authors:** Sourav Dey, Swati Shree Padhi, Rashmi Ranjan Behera, Sangita Lala, MM Arshida, Priyanka Das, Ruthrotha Selvi Bharathavikru

## Abstract

Altered cellular changes in tumors are regulated by a plethora of mechanisms, including post-transcriptional processes such as the reversible m6A RNA methylation which is shown to be involved in multiple steps of RNA processing thus playing a crucial role in gene expression. The individual roles of the writers, erasers and readers is slowly being unravelled and due to their regulatory role at the post transcriptional level, there are reports of both positive and negative modulatory effects on tumorigenesis. In this study, we have studied the role of one of the m6A writer complex proteins, WTAP and the m6A RNA demethylase FTO (fat mass and obesity associated protein) in 2D and 3D culture systems of triple negative breast cancer cells. Genome edited cell lines where WTAP or FTO levels were downregulated show altered cell proliferation through m6A mediated regulation of cell cycle regulators, and altered cell migration properties through modulation of EMT markers. The 3D spheroidal cultures derived from these genome edited lines revealed a role for m6A regulation in CSC marker expression, thus suggesting a potential oncogenic feedback loop between m6A modulators and the major hallmarks of cancer.

## Introduction

Eukaryotic gene expression is regulated at various steps including RNA Processing. RNA metabolism and its fate is regulated through chemical modifications on RNA. The most common RNA modification is methylation at the N6 position of the adenosine residue, referred to as m6A, which is known to regulate key cellular events such as, differentiation (**1**), proliferation and migration (**2**), metabolism (**3**), autophagy (**4**), senescence associated DNA Damage Responses (**5**), and cell death (**6**). The m6A modification is catalyzed by the methyl transferases which recognize the consensus RRACH motif, in association with other proteins, together referred to as the writer complex. The modification is reversible and removed by the demethylases, or erasers which are FTO and ALKBH5 which work through the 2-oxoglutarate dependent mechanism (**7, 8, 9**). The m6A epi-mark is decoded by the reader family of proteins including YTHDF and YTHDC family of proteins, deciding the fate of the mRNA in physiology and disease. Thus, the m6A modification plays a crucial role in several biological processes, misregulation of which can lead to human malignancies (**10, 11**).

Several cancers have shown altered activity of m6A readers, writers and erasers (**12**). The WMM (WTAP- METTL3/METTL14) complex is the m6A writer enzyme complex, where METTL3/METTL14 are the main catalytic enzymes, and WTAP recruits them to the RNA in nuclear speckles (**13**). WTAP is reported as an oncogene in most cancers in both m6A RNA methylation-dependent and independent manner. Abnormal expression of WTAP is associated with the progression of several types of cancers, including those of the brain (**14**), liver (**15**), oesophageal (**16**), lung (**17**), leukemia (**18**), lymphoma (**19**), and others. WTAP contributes to cancer progression by altering the RNA methylation, cell cycle, cell proliferation, cell growth, apoptosis, autophagy, EMT, and other biological functions (**20, 21**). The m6A RNA demethylases (FTO-Fat mass obesity and ALKBH5) have also been shown to be associated with cancer progression by contributing to metastasis, CSC maintenance and differentiation (**22, 23**). The demethylases remove the m6A by oxidative demethylation processes. FTO catalyses m6A conversion into N6- hydroxymethyl adenosine (hm6A) and gradually releases adenosine (A) and formaldehyde. In contrast, ALKBH5 releases formaldehyde rapidly by direct conversion of m6A to A (**24**). The RNA demethylase FTO, loss of function of which leads to severe growth defects, (**25, 26**) contributes to tumorigenesis through its regulatory roles on cellular proliferation (**27, 28, 29**). FTO regulates various cell cycle genes such as *Cyclin-D1* and thus contributes to the tumor phenotype (**29**). The RNA demethylase ALKBH5 has been shown to regulate Leukaemia stem cell pools as well as breast cancer stem cell population under hypoxic conditions. (**30**).

TNBC, Triple-negative breast cancer, where Estrogen receptor, Progesterone receptor, and HER2 are not expressed, is more aggressive among Breast Cancer. Altered expression of the WMM complex is associated with breast cancer tumorigenesis, where WTAP and METTL3/14 have been reported as oncogenic in TNBC. It is reported that METTL3 upregulates the oncoprotein FAM83D through m6A modification, contributing to tumorigenesis (**31**). WTAP has been shown to promote breast cancer progression through enhanced m6A deposition and expression of Enolase-1 (**32**), as well as through the WTAP/DLGAP1- AS1/miR-299-3p axis (**33**). lncRNA MALAT1 dependent stabilization of WTAP, leads to a tumor hypoxic environment thus contributing to TNBC progression through altered EMT markers (**34**) In TNBC, tumor size and grade were favorably correlated with WTAP expression (**35**). WTAP has also been shown to regulate alternative splicing and cell cycle in other cell types (**13**) however, the interplay of the various regulatory roles of WTAP and their contribution to TNBC progression is still unclear.

FTO-mediated demethylation downregulates the tumor suppressor BNIP3 in breast tumors (**36**). FTO polymorphism was identified in Her-2 negative breast cancer patients making it a potential therapeutic target (**37**). FTO has also been shown to influence the energy metabolism in breast cancer, thus contributing to cancer progression and tumorigenesis (**27**). In TNBC, FTO acts as a tumor suppressor, where it modulates the m6A dependent maturation of miRNA miR-17-5P and blocks its downregulation of tumour suppressor, ZBTB4 (**38**). The m6A demethylase, ALKBH5, has also been reported as a dual regulator in various cancers. In TNBC, ALKBH5 has been shown to reduce the m6A level of FOXO1A mRNA, thus promoting cancer (**39**). ALKBH5 is reported to upregulate UBE2C expression by altering m6A level and decreased p53 expression, which in turn contributed to the stemness, proliferation, and metastasis of the TNBC cells (**40**). The TNBC aggressive property is maintained by the m6A-dependent cell growth and survival, where the reader YTHDF2 also plays an essential role (**41**). YTHDC1-dependent m6A has been shown to regulate SMAD3 expression, and nuclear export thus enhancing the TGF-β signaling cascade in TNBC metastasis (**42**). TNBC has been shown to have high proliferative, metastasis, tumor recurrence. TNBC exhibits a higher proportion of Cancer Stem Cells (CSC) among breast cancer subtypes, contributing to its aggressive behaviour (**43, 44**).

Cancer stem cells (CSC) are the small subset of cells in the tumor bulk having self-renewal and chemo resistant properties, making it an ideal therapeutic target. There is growing evidence that mRNA modification is instrumental in the lineage specific differentiation and cellular reprogramming, which is altered during oncogenic transformation. Recent studies have shown that in most aggressive cancers, the key driver of intra tumor cancer heterogeneity are cancer stem cells (CSCs) or Tumour initiating cells (TICs) (**45, 46, 47, 48, 49, 50**). CSCs can originate from tissue resident adult stem cells, differentiated tissue cells, tumour cells, and tumour supporting cells. Identifying and characterizing CSCs are important for understanding the origin of CSC during cancer development and cancer relapsing (**51, 52**). Although there are several studies on the role of signalling pathways, cell surface markers and transcription factors in CSC maintenance (**53**), there are very few reports that have investigated the role of the RNA modification on the CSCs.

The role of the m6A modifiers in cancer has shown contradicting evidence, especially the involvement of WMM and erasers and thereby the overall impact of m6A modification. We decided to investigate this using a comparative knockout approach where both a writer complex protein as well as an eraser protein were depleted in the TNBC cells, to study their respective role on different cellular properties. The KO of the methyltransferase METTL3 has been shown to alter m6A levels, however we decided to focus on the RNA targets that are modulated by WTAP because of its additional involvement in alternative splicing (**54, 55**). Alternative splicing is a characteristic of aggressive tumours that promotes EMT. Thus investigating the regulatory role of WTAP, will allow us to study the interplay of various RNA processing events in the process of tumorigenesis. Along with this, we looked at FTO to validate the observations since changes modulated by m6A modification should demonstrate an opposite effect in the cells that lack FTO. WTAP deleted cells (WTAP KD) showed a decrease in cell proliferation and viability whereas the FTO deleted cells (FTO KD) showed an accelerated cell proliferation suggesting the role of m6A modulation on cell proliferation. Previously, the role of cell cycle regulators such as *Cyclin-D1* and *Cyclin-A2* have been shown to be regulated by the m6A modification (**56, 57**). Our results validated the findings that m6A modifications regulate *Cyclin-D1* expression, indicated by its downregulation in the WTAP KD cells whereas it was found to have upregulated expression in the FTO KD cells. We resorted to the 3D spheroid model based CSC enrichment, where we observed that WTAP KD spheroids were significantly smaller and FTO KD spheroids were larger in size, implying a role for WTAP in CSC maintenance. In contrast, FTO represses CSC formation, thus highlighting a role for m6A modulation of the CSC phenotype. Since 30% of the TNBC patients have been shown to be associated with metastasis, we tested cell migration in these knockdown cells. Cell migration potential of WTAP KD cells was lower as compared to FTO KD cells, suggesting a role for m6A modification in the Epithelial Mesenchymal Transition and thus in tumor progression. Taking all these observations together, we conclude that the m6A modification is a regulator of cell proliferation, CSC maintenance, and cell migration thus contributing to aggressive tumour phenotype. This study thus highlights the importance of m6A modifier regulated mechanistic implications in tumour biology especially for TNBC and provides compelling evidence for exploring the m6A modification pathway for therapeutic targeting in TNBC.

## Results

### RNA Methylation modulators are differentially expressed in Triple Negative Breast Cancer Cells

We used the Kaplin-Meier (KM) plotter, to analyse the the m6A writer complex associated protein WTAP levels and its role in cancer, where we observed that it showed an oncogenic role, corresponding to a lower survival rate when WTAP expression was upregulated (**Fig 1A**). This was completely opposite in the case of the RNA demethylase, FTO, which showed minimal variation and a correlation with better survival when FTO was highly expressed (**Fig 1B**). However, when a similar analysis was done for ALKBH5, no significant correlation was observed (**Fig. S1A)**, suggesting an important role of WTAP in cancer progression. Analysis of publicly available TCGA datasets, revealed that while WTAP expression was consistently upregulated in most breast cancer patients, the demethylase FTO showed varied expression and was mostly downregulated (**Fig 1C)**. To characterize the cellular and molecular changes that are associated with the altered expression of RNA methylation modulators, WTAP and FTO KDs were created in the Triple Negative Breast Cancer (TNBC), MDA-MB-468 cell line (**Fig S1B**). Two different guide RNAs were used for WTAP and FTO, as detailed in **Figure S1C and S1D** respectively, and the cells were subjected to puromycin selection. The WTAP and FTO knockdown was confirmed through Immunofluorescence analysis (**Fig. S1E and S1F, Panel I**) where images were captured and quantified as represented in Panel II (**Fig. S1E and S1F** respectively). Further, Western blot analysis showed that in comparison to the corresponding empty vector (EV) control more than 50% of WTAP expression was decreased in the MDA-MB-468 cell line (henceforth referred to as W-EV, W-KD1 and W-KD2), (**Fig 1D, Panel I**, and quantified (**Panel II)**. Nearly 50% FTO expression was decreased in the MDA-MB-468 cell line (henceforth referred to as F-EV, F-KD1, F-KD2), (**Fig 1E, Panel I**), and quantified in **Panel II**.

**Figure 1:**
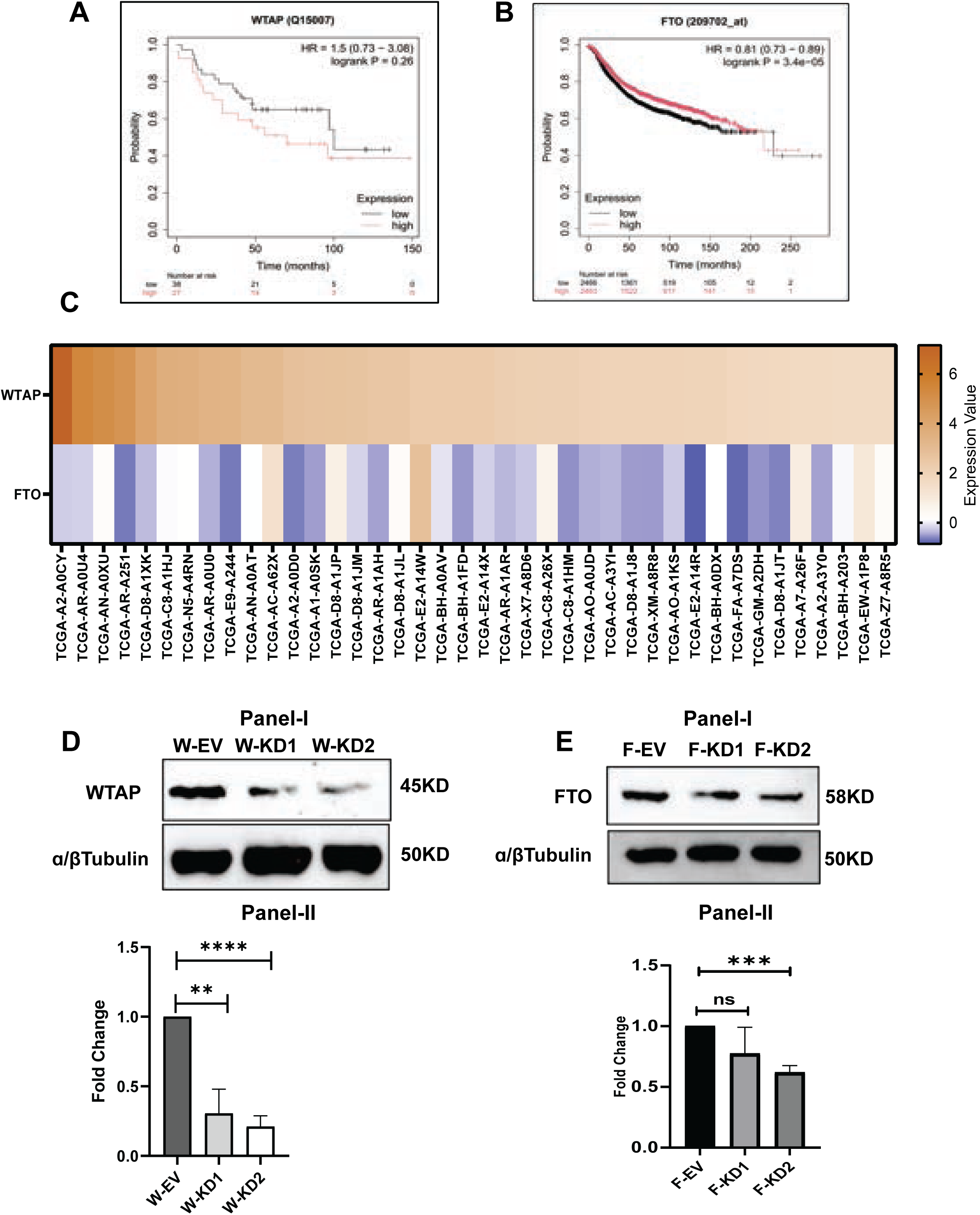
RNA Methylation modulators are differentially expressed in Triple Negative Breast Cancer Cells. **A.** Kaplan-Meier Plotter generated survival curves of breast cancer patients in WTAP high and low expression conditions. **B.** Kaplan-Meier Plotter generated survival curves of breast cancer patients in FTO high and low expression conditions. **C**. Heatmap of GDC data portal generated gene expression of WTAP and FTO genes from 40 breast cancer patient samples of TCGA dataset. Z-score transformed gene expression values are plotted. **D.** Western Blotting image of WTAP KD in MDA-MB-468 cells, (**Panel-I**). Graphical representation of fold expression of WTAP in knockdowns (W-KD1, W-KD2) with respect to empty vector control (W-EV), normalized to housekeeping control ⍺/β-Tubulin, calculated by ImageJ analysis (**Panel-II**). The error bars depict the Means and SD, n=3, Significance analysis performed using student’s t-test by GraphPad, P-Value< 0.01 =**, <0.0001=****. **E**. Western Blotting image of FTO KD in MDA-MB-468 cells (**Panel-I**). Graphical representation of fold expression of FTO in knockdowns (F-KD1, F-KD2) with respect to empty vector control (F-EV), normalized to housekeeping control ⍺/β-Tubulin, calculated by ImageJ analysis (**Panel-II**). The error bars depict the Means and SD, n=3, Significance analysis performed using student’s t-test by GraphPad, P-Value <0.001 =***.

### m6A Modifiers, WTAP and FTO Regulate Cell Proliferation in Triple Negative Breast Cancer Cells

m6A modification and the WMM complex protein, WTAP, have been shown to regulate cell cycle changes. Hence, we decided to investigate the effect of knockdown of m6A modulators on various parameters including cell viability and cell proliferation (**Fig. S2A**). The WTAP KD cells when tested for cell viability using the MTT assay, showed a 50% decrease in cell viability compared to the control (**Fig. 2A)**. This observation was further corroborated with a cell proliferation analysis where WTAP KD cells were observed to show a significant decrease in cell count in 24 hours compared to the EV control (**Fig. S2B**). The proliferative index of WTAP KD cells was measured using Immunofluorescence for Ki67 expression (**Fig 2B, Panel I**) and quantified as represented in panel II, which showed a more than 50% decrease in cell proliferation. To observe the effects of FTO KD on cell viability, we performed an MTT assay on the knockdown cell line, which showed a significant increase in cell viability as compared to the empty vector control (**Fig. 2C**). We further checked the proliferation status of FTO KD cell lines and observed an increased cell number in the knockdown cells after 24 hours of proliferation as compared to empty vector control (**Fig S2C**). Proliferation index of FTO KD cells was further analyzed by immunofluorescence assay using Ki67 antibody, which displayed a significantly increased fluorescence as compared to the empty vector control (**Fig. 2D, Panel I, quantification in Panel II**), suggesting that any alteration in the expression of the m6A modulators, directly influences cell viability and cell proliferation.

**Figure 2:**
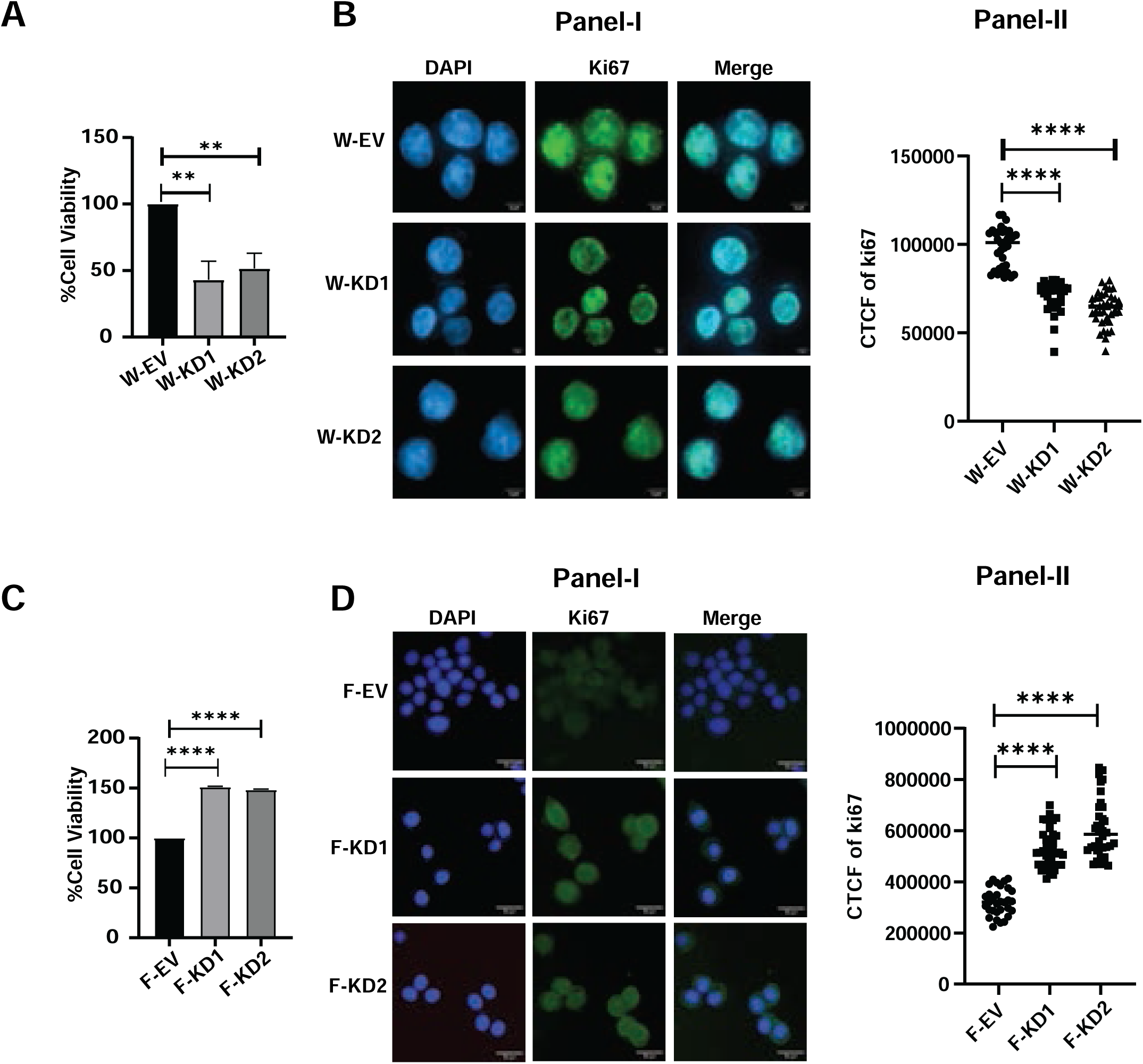
m6A Modifiers, WTAP and FTO Regulate Cell Proliferation in Triple Negative Breast Cancer Cells. **A.** Graphical representation of percentage of cell viability in WTAP KD cells (W-KD1, W-KD2), compared to empty vector control (W-EV) measured by MTT Assay. The error bars depict the SD and Mean, n=3, Significance analysis using student’s t-test performed by GraphPad, P-Value< 0.01 =*\*\**. **B.** Immunostaining image of Ki67 (Green Channel), DAPI (blue channel) and merged image of WTAP KD cells (**Panel I**). Graphical representation of the corrected total fluorescence intensity calculated by ImageJ analysis (**Panel-II**). The error bars depict the SD and Mean, n=3, Significance analysis using student’s t-test performed by GraphPad, P-Value<0.0001=****. **C**. Graphical representation of percentage of cell viability in FTO KD cells (F-KD1, F-KD2), compared to empty vector control (F-EV) measured by MTT Assay. The error bars depict the SD and Mean, n=3, Significance analysis using student’s t-test performed by GraphPad, P-Value<0.0001=****. **D.** The immunostaining image of Ki67 (Green Channel), DAPI (Blue Channel) and merged image of FTO KD cells (**Panel-I**). Graphical representation of the corrected total fluorescence intensity calculated by ImageJ analysis (**Panel-II**). The error bars depict the SD and Mean, n=3, Significance analysis using student’s t-test performed by GraphPad, P-Value<0.0001=****.

### *Cyclin* expression is regulated in an m6A dependent manner in the TNBC cells

The altered cell proliferation parameters as shown above, indicate a potential role of altered cell cycle regulators. To understand if these changes are reflected at individual cell cycle stages, we performed a FACS analysis using Propidium Iodide staining. The cell cycle stages for both WTAP KD (**Fig. S3A)** and FTO KD (**Fig. S3B**) cells are represented in a histogram. WTAP downregulation led to an increased accumulation of cells in the G0/G1 stage in WTAP KD (W-KD1, W-KD2) cells compared to empty vector control, W-EV (**Fig 3A**) which strengthened the earlier observation of decreased cell viability. In the FTO KD cells (F-KD1, F-KD2), decrease in the G1 cells and an increased accumulation of cells in the G2/M stage compared to empty vector control cells (F-EV), was observed (**Fig 3B**). Earlier studies have shown that WTAP modulates *Cyclin-D1* expression and FTO regulates *Cyclin-A2* expression (**57**). We therefore assessed the expression of *Cyclins* using qRT PCR and observed a downregulation of *Cyclin-D1* and *Cyclin-A2* expression in the WTAP KD cells (W-KD1, W-KD2) as compared to the empty vector controls, W-EV (**Fig 3C**). In-contrast, the FTO KD cells (F-KD1, F-KD2) showed an upregulation of *Cyclin-D1* and *Cyclin-A2* expression as compared to empty vector control, F-EV (**Fig 3D**). To confirm whether this modulation of the Cyclin expression is mediated through m6A levels, we performed an m6A RNA Immunoprecipitation (RIP) followed by qRT PCR (**Fig. S3C**) and validated the same using primers for known m6A targets (**36**) (**Fig. S3D**). The m6A RIP showed a decrease in m6A enrichment on *Cyclin-D1* and *Cyclin-A2* 3’ UTR in the WTAP KD cells (W-KD1, W-KD2), compared to empty vector control W-EV (**Fig 3E**). However, in the FTO KD cells (F-KD1 F-KD2), an increased enrichment of m6A on the *Cyclin D1* was observed, compared to the empty vector control, F-EV (**Fig. 3F)**. Taken together, these results provide evidence for an m6A dependent regulation of cell cycle change in the TNBC cell lines, strengthening the conclusion that the m6A modulators have an important regulatory role in tumorigenesis.

**Figure 3:**
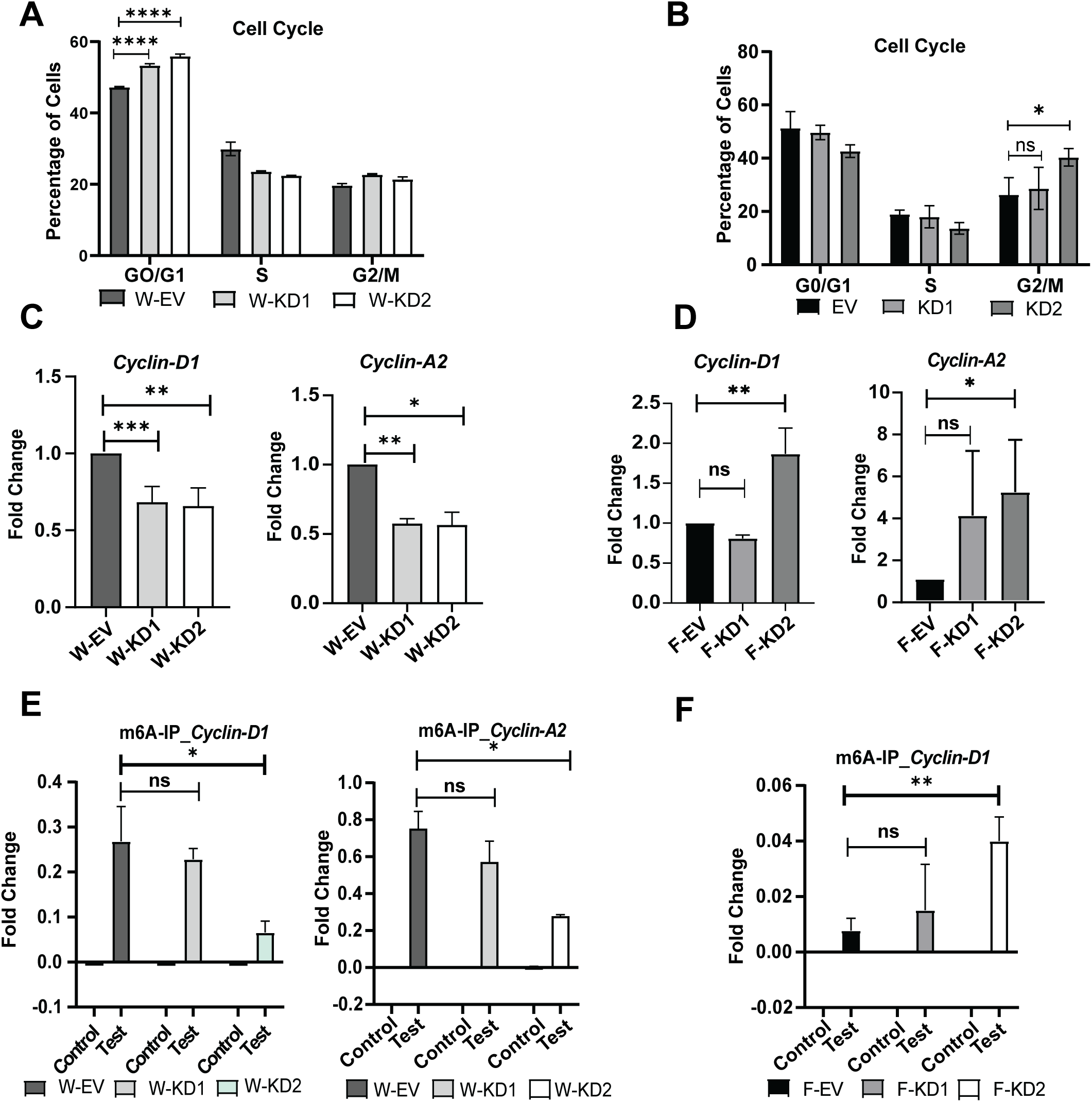
*Cyclin* expression is regulated in an m6A dependent manner in the TNBC cells. **A.** Graphical representation of flow cytometry-based analysis of cell cycle stages G0/G1, S, and G2/M for WTAP KD (W-KD1, W-KD2) and EV control (W-EV). Percentages of cells from each stage are represented graphically for both the Control and knockdown cells. The error bars depict the SD and Mean, n=3, Significance analysis using student’s t-test performed by GraphPad, P-Value<0.0001=****. **B.** Graphical representation of flow cytometry-based analysis of cell cycle stages G0/G1, S, and G2/M for FTO KD (F-KD1, F-KD2) and EV control (F-EV). Percentages of cells from each stage are represented graphically for both the control and knockdown cells. The error bars depict the SD and Mean, n=3, Significance analysis using student’s t-test performed by GraphPad, P-Value< 0.05=*. **C.** qPCR analysis of cell cycle regulator genes *Cyclin-D1* and *Cyclin-A2* in WTAP KD cells (W-KD1 W-KD2) compared to W-EV, normalised with *GAPDH* housekeeping control. The error bars depict the SD and Mean, n=3, Significance analysis using student’s t-test performed by GraphPad, P-Value < 0.05=*, < 0.01 =*\*\**, <0.001 =***. **D.** qPCR analysis of cell cycle regulator genes *Cyclin-D1* and *Cyclin-A2* in FTO KD cells (F-KD1 F-KD2) compared to F-EV, normalised with *GAPDH* housekeeping control.The error bars depict the SD and Mean, n=3, Significance analysis using student’s t-test performed by GraphPad, P-Value < 0.05=*, < 0.01 =*\*\**. **E.** qPCR analysis of m6A enrichment of *Cyclin-D1* and *Cyclin-A2* 3’UTR in m6A-Immunoprecipitated (m6A-IP) RNA from WTAP KD (W-KD1, W-KD2) and EV control (W-EV). Expression was first normalised with input control and then fold expression was calculated in control-IP vs. Test-IP (m6A Antibody) and represented graphically. The error bars depict the SD and Mean, n=2, Significance analysis using student’s t-test performed by GraphPad, P-Value < 0.05=*. **F.** qPCR analysis of m6A enrichment of *Cyclin-D1* in m6A-Immunoprecipitated (m6A-IP) RNA from FTO KD (F-KD1, F-KD2) and EV control (F-EV). Expression was first normalised with input control and then fold expression was calculated in control-IP vs. Test-IP (m6A Antibody) and represented graphically. The error bars depict the SD and Mean, n=2, Significance analysis using student’s t-test performed by GraphPad, P-Value<0.01 =*\*\**.

### RNA Methylation modulates Cancer Stem Cell Maintenance in 3D spheroidal cultures of TNBC

The role of m6A modification in stem cell fate commitment has been shown to contribute to tumorigenesis (**58**). Since the experiments were all done in a TNBC cell line model, we were interested in knowing whether the m6A modification might have a role in cancer stem cell maintenance. Since the spheroidal cultures can be used to enrich for CSCs (**59, 60**), we created mammospheres using the WTAP and FTO KD MDA-MB-468 cell lines (**Fig. S4A**). We observed that the W-KD1 and W-KD2 lines had reduced spheroid forming ability as compared to the EV control during a 7 day spheroid formation assay. Day-5 and day-7 spheroids of WTAP KD were smaller compared to W-EV (**Fig. 4A**). The Image-J based analysis of spheroid size and number of day-5 and day-7 WTAP KD spheroids compared to W-EV, showed a significant decrease (**Fig. 4B, panel-I, panel-II** respectively). Contrastingly, the day-5 and day-7 spheroids of FTO KDs were larger compared to F-EV (**Fig. 4C**). In case of FTO, the Image-J based analysis of spheroid size and number showed an increase in FTO KDs compared to F-EV (**Fig. 4D, panel-I, panel-II** respectively). Since the spheroid formation ability corroborated the results observed in cell proliferation analysis, we further went ahead to investigate the *Cyclin* levels in the spheroids of the genome edited cell lines. We observed a net decrease of both *Cyclin-D1* and *Cyclin-A2* in W-KD1 and W-KD2 cell lines as compared to W-EV (**Fig. S4B, Panel I, Panel II respectively).** The FTO-KD spheroids, were checked for cell proliferation using, immunostaining for Ki67, which showed an overexpression of the proliferative marker Ki67 in F-KD1 and F-KD2 spheroids compared to F-EV spheroids (**Fig S4C**). The spheroidal culture was checked for the CSC marker expression using qRT-PCR analysis. Downregulation of CSC markers *CD133, SOX2, KLF4* expression was observed in the W-KD1 and W-KD2 cells compared to W-EV (**Fig. 4E**), whereas the F-KD1 and F-KD2 spheroids showed an upregulation of the CSC marker *CD133, SOX2* and *KLF4* expression in comparison to the corresponding EV control (**Fig. 4F**). Additionally, WTAP KD spheroids were immunostained to look for SOX2 expression which also showed decreased expression of SOX2 in W-KD1 and W-KD1 spheroids compared to W-EV spheroids (**Fig S4D).** Taken together, these observations clearly show a role for the m6A modulators in regulating the CSC niche and thus contribute to tumorigenesis.

**Figure 4:**
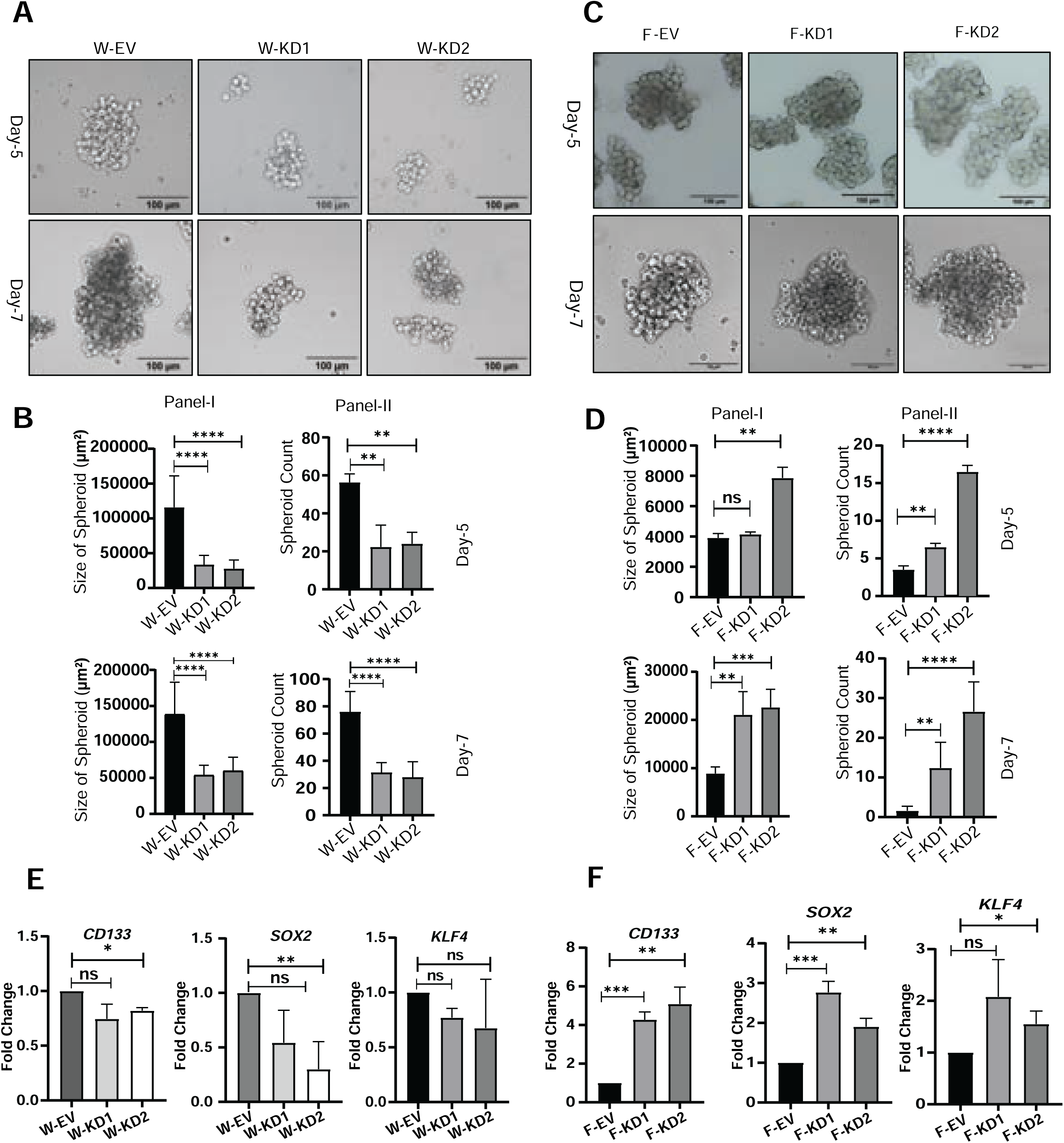
RNA Methylation modulates Cancer Stem Cell Maintenance in 3D spheroidal cultures of TNBC. **A.** Brightfield images of Day-5 and Day-7 spheroid growth of WTAP KD (W-KD1, W-KD2) and empty vector control (W-EV). **B.** Graphical representation of Size (**Panel-I**) and number (**Panel-II**) of Day-5 and Day-7 spheroids of WTAP KD (W-KD1, W-KD2) and empty vector control (W-EV) calculated by ImageJ analysis. The error bars depict the SD and Mean, n=3, Significance analysis using student’s t-test performed by GraphPad, P-Value< 0.05=*, < 0.01 =*\*\**, <0.0001=****. **C.** Brightfield images of Day-5 and Day-7 spheroid growth of FTO KD (F-KD1, F-KD2) and empty vector control (F-EV). **D.** Graphical representation of Size (**Panel-I**) and number (**Panel-II**) of Day-5 and Day-7 spheroids of FTO KD (F-KD1, F-KD2) and empty vector control (F-EV) calculated by ImageJ analysis. The error bars depict the SD and Mean, n=3, Significance analysis using student’s t-test performed by GraphPad, P-Value< 0.01 =*\*\**, <0.001 =***, <0.0001=****. **E.** qPCR analysis of cancer stem cell markers, *CD133*, *SOX2*, and *KLF4* in WTAP KD spheroids compared to EV control spheroids (W-EV), normalised with housekeeping control *GAPDH*. The error bars depict the SD and Mean, n=3, Significance analysis using student’s t-test performed by GraphPad, P-Value < 0.05=*, < 0.01 =*\*\**. **F.** qPCR analysis of cancer stem cell marker genes *CD133*, *SOX2*, and *KLF4* in FTO KD spheroids compared to EV control spheroids (F-EV), normalised with housekeeping control *GAPDH*. The error bars depict the Means and SD, n=3 Significance analysis using student’s t-test performed by GraphPad, P-Value < 0.05=*, < 0.01 =*\*\**, <0.001 =***.

### Epithelial Mesenchymal Transition associated markers and the alternative splicing contributes to tumorigenic potential of TNBC

The m6A modulators regulate the CSC markers in TNBC derived spheroid models as seen in **Fig.4E** and **4F.** *CD44* is one such CSC marker which is known to regulate EMT in TNBC through isoform specific expression. *CD44* exists as three different isoforms *CD44S* (Standard), *CD44T* (Total), and *CD44V* (Variant). *CD44V*, overexpression has been observed in several cancers and contributes to the aggressiveness of cancer by regulation of oncogenic signalling pathways, cell migration, metastasis and chemoresistance (**61**). We analyzed the expression of *CD44* in WTAP KD spheroids where W-KD1 and W-KD2 showed a downregulation in knockdowns as compared to the empty vector control, W-EV (**Fig. S5A**). The isoform specific *CD44* expression (*CD44S, CD44T, CD44V*) was assessed through qRT PCR using isoform specific primers, which showed a downregulated expression in WTAP KD spheroids compared to empty vector, W-EV spheroids (**Fig. 5A**). The relative fold change of the two isoforms *CD44S* and *CV44V* was calculated over the *CD44T* isoform, which revealed that the *CD44V* expression was downregulated in the knockdowns in comparison to the *CD44S* expression level **(Fig. 5B**). This observation suggested the role of the m6A modifier WTAP in isoform switching from *CD44S* to *CD44V*. Since the *CD44V* isoform also regulates EMT and metastasis in breast cancer (**62**), we decided to assess the expression of the EMT markers, including the Mesenchymal markers, *SNAI1 VIM, N-Cadherin, TWIST1* and the epithelial markers, *FN1, E-Cadherin*, from previously reported TCGA datasets on TNBC samples. Upon analysis of these datasets, higher WTAP expression was found to be associated with high expression of EMT markers, especially *VIM and SNAI1* and correspondingly, low expression of epithelial marker *E- cad,* and the converse with higher WTAP expression (**Fig. 5C**). The expression pattern of all the above EMT markers when analyzed with respect to FTO expression, showed the reverse to what was observed with WTAP (**Fig. S5B**). EMT marker, *TWIST1* expression was analysed in the WTAP downregulated TNBC cells where W-KD1 and W-KD2 showed downregulation compared to the empty vector control (W-EV) implying a role for m6A modulator regulated regulation of EMT (**Fig. 5D**).

**Figure 5:**
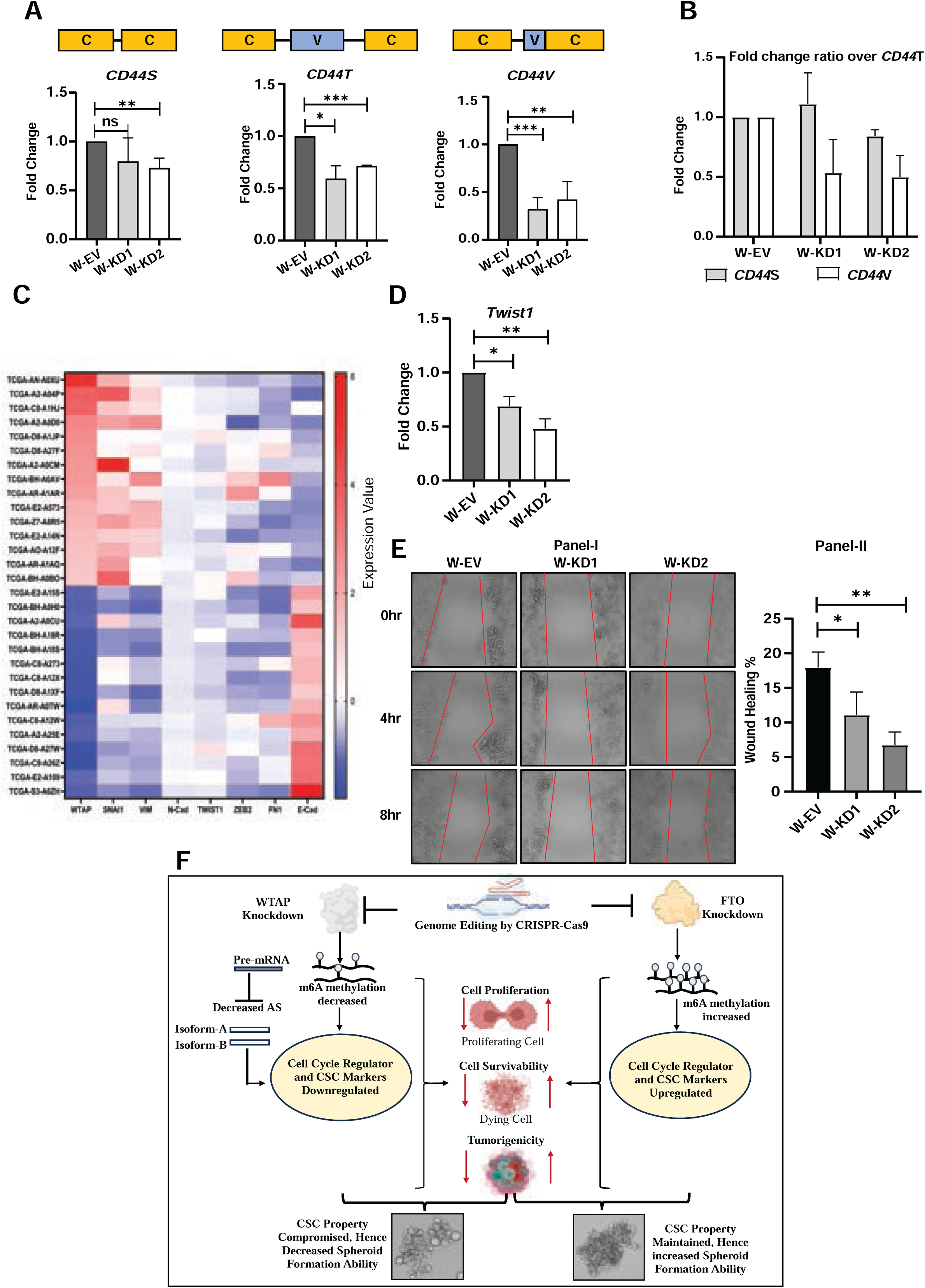
Epithelial Mesenchymal Transition associated markers and their alternative splicing contributes to tumorigenic potential of TNBC. **A.** qPCR analysis representing fold change expression of the three splice variants of *CD44, CD44T* (total), *CD44S* (standard) and *CD44V* (variable) in W-KD1 and W-KD2 cells compared to W-EV control, normalised with housekeeping control *GAPDH* expression.. The error bars depict the SD and Mean, n=3, Significance analysis using student’s t-test performed by GraphPad, P-Value < 0.05=*, < 0.01 =*\*\**, <0.001 =***. **B.** Graphical representation of the fold change ratio of each isoform (*CD44S* and *CD44V*) over *CD44T* C. Heatmap of GDC data portal generated gene expression of EMT markers (*SNAI1, VIM, N-cadherin, TWIST1, ZEB2, FN1 and E-cadherin*) arranged in ascending order of WTAP expression. Z-score transformed gene expression values are plotted. **D.** qPCR analysis of EMT marker gene expression (*TWIST1*) in WTAP KD TNBC cells (W-KD1, W-KD2) compared to EV control (W-EV), normalised with housekeeping control *GAPDH* expression. The error bars depict the SD and Mean, n=3, Significance analysis using student’s t-test performed by GraphPad, P-Value< 0.05=*, < 0.01 =*\*\**. **E.** Brightfield image of wound healing (cell migration) at different time points (0 hour, 4 hours, and 8 hours) in WTAP KD (W-KD1, W-KD2) cells compared to EV control, W-EV (**Panel-I**). The percentage of cell migration was calculated by ImageJ and represented graphically (**Panel-II**). The error bars depict the Means and SD, n=3, Significance analysis using student t-test performed by GraphPad, P-Value < 0.05=*, < 0.01 =*\*\**. **F.** Schematic showing the role of m6A modifiers (WTAP and FTO) dependent regulation in TNBC.

Since EMT is a hallmark associated with cell migration and tumorigenesis, we decided to investigate the role of WTAP in a wound healing assay. The WTAP KD cells showed a decreased cell migration ability compared to the W-EV cells, during 8 hours of migration analysis (**Fig. 5E, Panel I**), The percentage cell migration represented in **Panel II**, showed nearly 15% less migration ability of W-KD1 and W-KD2 compared to W-EV. The FTO KDs cell on other hand showed an increased migration ability in comparison to F-EV (**Fig. S5C, Panel-I**), which is about 10% increased in F-KD1 and F-KD2 compared to F-EV (**Fig. S5C, Panel-II**). Taken together, this observation proves a role of the m6A modifiers in cell migration by regulating EMT marker expression, and through alternative splicing mediated by WTAP. In conclusion, this study shows a role for the m6A writer complex protein, WTAP in regulating cell cycle through m6A modification, modulating cancer stem cell enrichment through surface marker gene expression, alternative splicing as well as EMT marker expression, thus contributing to tumorigenesis (**Fig. 5F**). Whereas the m6A writer complex protein, WTAP seems to contribute to tumorigenicity in TNBC, the m6A RNA demethylase, FTO shows a tumour suppressive role, thus providing evidence for the m6A RNA methylation dynamics to be an important contributing factor in RNA processing mediated mechanism contributing to TNBC manifestation and progression.

## Discussion

In this study, we have explored the role of m6A modifiers, WTAP and FTO in regulating the m6A modification in TNBC cancer stem cell maintenance. This regulation is a multi- step process that involves m6A mediated regulation of cell cycle regulators such as *Cyclin-D1* and *Cyclin-A2*, which directly influences cell proliferation in both monolayer and spheroidal culture systems. The m6A modifiers also contribute to the overall tumorigenic potential of TNBC by regulating the expression of cancer stem cell marker genes and EMT associated marker genes, thus providing a niche that supports the aggressive nature of TNBC.

The dynamicity of the m6A mark is maintained by the interplay of the methylation writer complex and the erasers. Although the WMM complex (WTAP, METTL3/14), functions as the writer complex (**13**), WTAP has been singled out to be oncogenic in most cancers (**20**), suggesting that WTAP might have roles other than those that are mediated by the m6A modification itself. Incidentally, WTAP has been shown to have a role in alternative splicing and cell cycle regulation (**63**). The eraser proteins, FTO and ALKBH5 have been shown to be associated with different cancers, albeit with contradictory roles (**64**). In this study, we observe that in breast cancer samples, both the m6A writer complex protein WTAP and Erasers show altered expression, which also corresponded to their association with survival, as seen by KM plots, although ALKBH5 expression did not show correlation with survivability. Among breast cancer, TNBC is the most common and aggressive cancer, and no studies have assessed the regulatory role of m6A dynamicity in TNBC, which motivated us to use MDA-MB-468 cell lines as the exemplar cell line model to investigate the role of m6A dynamicity in TNBC tumour manifestation and progression.

The genome edited lines of the m6A modulators showed altered cell proliferation rates. Between the replicates there was change in the proliferation index, which could reflect the variation in the efficiency of the knockdown itself between the two biological replicates. The cell cycle analysis showed accumulation of cells in G0/G1 stage in WTAP KD cells whereas the FTO KD cells showed accumulation in G2/M stage. Incidentally, it has been shown that WTAP regulates the G2/M stage of cell cycle by stabilizing *Cyclin-A2* (**62**) and alters the G0/G1 stage by regulating the cell cycle regulator *Cyclin-D1* (**65**). We now show that in the TNBC cells, WTAP plays a role in the cell cycle entry point at the G0/G1 stage itself. However, the analysis using the FTO genome edited lines shows a role for m6A modulation even at the G2/M stage. Thus, the m6A modification seems to regulate cell cycle through regulation of multiple cell cycle regulators. *Cyclin-D1* activates the CDKs that are involved in phosphorylation of RB proteins leading to degradation which allow the E2F factors to facilitate G1 to S phase specific gene expression (**66**). *Cyclin–A2*, a crucial regulator of the G2/M phase of transition, binds to *Cdk1* and activates it, which phosphorylates the proteins for M-phase transition. *Cyclin-A2* also promotes the DNA synthesis in S-phase (**66, 68**). Stabilization and overexpression of these regulators through WTAP or FTO mediated modulation, thus leads to altered cell cycle stages.

We have observed the m6A dependent cell cycle alteration and proliferation contributes to the cancer progression in TNBC. TNBC is characterized by aggressive growth, continuous proliferation (**69**), invasiveness, metastasis (**70**), heterogeneity and chemo-resistance (**71, 72**), thus hinting at a role of cancer stem cells in the tumorigenicity. Earlier studies have shown the enrichment of breast cancer stem cells (BCSC) in TNBC, as one of the reasons for aggressiveness and poor prognosis (**73**). Research has shown the regulatory role of m6A in BCSC, on self renewal, resistance to therapy, tumorigenesis and differentiation (**74, 75**), we decided to study the role of the m6A modulators especially WTAP and FTO in BCSC. It has been also reported that the presence of a high proportion of cancer stem cells in TNBC increases the sphere formation ability (**76**). We used the 3D spheroidal model to understand the role of the m6A modifiers in the BCSC population. Interestingly, we observed a decreased spheroid formation efficiency both in terms of size and number in the WTAP KD cells whereas the FTO KD cells led to large well defined spheroids. The cell cycle directly affects the growth rate of spheroids. Cells in G2 and M phase are considered as more active proliferative stages that contribute to the spheroid growth. We have also observed the m6A modulated alteration of cell cycle regulators *Cyclin-A2* and *Cyclin-D1* mediated enhancement of spheroid formation. Our study was restricted to examining the spheroid ability till day 7, since the WTAP spheroids which were already smaller in size, started to disintegrate. However if tested using in vivo conditions with the tumor niche this mechanism can be studied clearly.

The maintenance of the CSC pool is regulated by not just the intrinsic mechanisms but also, the TME consisting of multiple immune cells, stromal cells, extracellular matrix, specific conditions like low pH and hypoxia, and the associated signaling cascades (**77, 78**). The CSC markers, especially *CD44* and its alternative splicing, are altered corresponding to the m6A modulators expression, thus confirming the change in CSC characteristics. Although the role of WTAP in alternative splicing has been studied in developmental context, the exact mechanism of its role in *CD44* isoform switching needs to be studied. Additionally, a detailed study that examines the individual contribution of the different TME components and the corresponding signalling pathways is required to chart out the exact molecular details associated with m6A modification in TNBC manifestation. The m6A modulator knockdown cells show an alteration in their cell migration ability within 8 hours as checked by the wound healing assay, which suggests that the invasiveness of the cells can be regulated by the m6A modification. Extending this study for a longer duration can lead to more than the observed 10-15% change in cell migration.

TNBC is considered to be one of the most aggressive and most difficult to treat amongst breast cancers. Very few therapeutic targets are reported till date for TNBC, among which mostly studied is the PARP signalling pathway. PARP responds to DNA damage and is overexpressed in cancer cells. In TNBC, targeted therapies against few signaling pathways have been reported. Among these, the EGF signaling pathway, which is frequently overexpressed through upregulation of its receptor, EGFR, plays a key role in regulating cell proliferation, invasion, and angiogenesis. Gefitinib, an EGFR inhibitor, has been shown to target TNBC. Additionally, inhibitors of the TGF-β and mTORC signaling pathways have also been explored as potential therapeutic strategies (**79**). However, due to limited mechanistic insights into the RNA modification regulated molecular pathways driving TNBC and the associated chemoresistance and metastasis, there are few studies that have investigated their therapeutic potential. Our study highlights a role for m6A modification in regulating cell proliferation, cell cycle and CSC pool as well as cell migration in TNBC. Our study not only provides the mechanistic details into TNBC manifestation and progression, but also provides a compelling reason to target the m6A modification specifically on target genes or alteration of the expression of the writers and erasers of m6A modification as a therapeutic strategy.

## Material and Methods

### Cell Culture

MDA-MB-468 triple negative breast cancer (TNBC) cell lines and HEK293T were cultured in Dulbecco’s Modified Eagle Medium (Catalog No.-11965092, Gibco) with 10% Fetal Bovine serum (Catalog No.- 16000044, Gibco) supplement and 1% Pen-strep solution (Catalog No.- 15140122, Gibco). Cells were grown in a humidified chamber with 5% CO2 at 37℃. Cells were cultured and tested for mycoplasma routinely.

### Knockdown creation by CRISPR-Cas9 genome editing

WTAP and FTO KD MDA-MB-468 cell lines were created by the use of CRISPR-Cas9 genome editing. Primers for gRNA targets against WTAP and FTO were designed using the webtool ChopChop. Primers were annealed at 37℃ for 30 minutes in a thermal cycler and used for ligation at 16℃ overnight with digested lenti-CRISPR-v2 vector (**80**). After confirmation by sequencing, guide RNA target cloned vectors were transfected with packaging vectors pVSVg (AddGene #8454) and psPAX2 (AddGene #12260) to the HEK293T cells for viral packaging. After 36 hours of transfection, the viral titer was collected and filtered by a 0.45 micron syringe filter. Viral titer was then transduced to the MDA-MB-468 cell line with 0.8µg/ml Polybrene to edit the target genes. 2µg/ml puromycin (Catalog-no.- CAS 58-58-2) was used for two weeks for selection. Cell lysate was prepared by using RIPA buffer and quantified by Bradford assay (Catalog No.- B6916) and the knockdowns were validated by Western Blotting.

### Western Blotting

Cell lysates were prepared using RIPA lysis buffer (Tris-Cl-50mM, pH=8.0, NaCl-150mM, EDTA-2mM, NP-40-1%, Sodium Deoxycholate-0.5% and Sodium Dodecyl sulphate-0.1%) with 1X of Protease Inhibitor Cocktail (cOmplete™, Mini, EDTA-free, Sigma Aldrich). Protein separation was performed by SDS-PAGE, and proteins were transferred to the PVDF membrane. Proteins were incubated with primary antibodies at 1:1000 ratio (**Table-S1**) for overnight at 4℃ followed by respective secondary antibody incubation. By applying ECL (substrate and enhancer at 1:1 ratio) to the membrane, bands were visualised in the BioRad imaging system and images were captured. The band intensities were quantified using ImageJ, and the fold expression was calculated by normalizing with the housekeeping control.

### Immunofluorescence

25000 cells were counted and seeded on poly-L-lysine coated coverslip. The cells were allowed to grow for 24 hours and were then fixed with 4% formaldehyde (Catalog No.-100496). The fixed cells were permeabilized by 0.1% Triton-X-100 (Catalog No.- X100-100ML). Followed by washing with PBST (PBS +0.1% Tween-20), cells were incubated with primary antibodies (at 1:250, **Table-S1**), overnight at 4℃. The coverslips were subsequently washed for non-specific primary antibody binding and then incubated with Secondary antibody tagged with Alexa Fluor-488 (1:500, **Table-S1**) and incubated for 1 hr at room temperature in the dark.The cells were washed, counterstained with DAPI, mounted and visualized using Zeiss fluorescence microscope. The images were captured with 20X magnification. The images were processed and signals were quantified in ImageJ software. The ROI for 40-50 cells were selected with a background ROI and then fluorescence intensity was recorded. The corrected fluorescence intensity was calculated for each cell and then plotted in a graph.

### MTT Cell Viability Assay

Cell viability was determined using MTT (3-(4,5-Dimethylthiazol-2-yl)-2,5-Diphenyltetrazolium Bromide, HiMedia Catalog No.- MB186-1G) cell viability assay. 5000-7000 cells were seeded in a 96-well cell culture plate and grown for 24 hours. Cells were washed with PBS and then treated with MTT reagent (working concentration 0.5 mg/ml) at 37°C for 4 hours. The Formazan crystals were dissolved by DMSO (HiMedia, GRM5856). The absorbance of the dissolved product was measured at 570 nm by UV plate reader. Absorbance was normalised with the blank control and then cell viability was calculated against Empty vector treated control. Each experiment was done in technical triplicate and the experiment was performed with three biological replicates.

### Wound Healing Assay

Wound healing of WTAP-KD and FTO KD MDA-MB-468 with control cells (W-EV and F-EV respectively) was performed. 50000 cells for each cell type were seeded in a 12 well plate and allowed to grow until 90% confluency. Then a scratch or wound was created using a 10µl sterile micropipette on the monolayer. The scratched cells were washed with PBS and were imaged at 0 hour. Cells were replenished with complete media and allowed to grow. Wound images were captured at 0 hour, 4 hour, 8 hour, using Zoe Fluorescent imager (Bio Rad) at 20x magnification. The cell movement within the scratches were analyzed and quantified using imageJ software. The wound area for each time point of cells (0 hour, 4 hour, 8 hour) for both knockdowns and controls was measured by the software and recorded. The percentage movement was calculated with reference to the 0 hour time point and plotted in a graph.

### Cell cycle analysis by Flow Cytometry

Cell cycle analysis was performed on WTAP and FTO KD MDA-MB-468 cells using Flow cytometric analysis. Cells were seeded at a density of 1 x 10^6^ cells in a six well plate. The cells were harvested and washed with PBS. Cells were subsequently fixed with 70% ethanol and incubated at -20℃ for 30 minutes. The cells were centrifuged at 3000 rpm for 5 min and washed with PBS. The cells were then treated with Propidium Iodide (PI) (Catalog No.-P4170) containing RNase and were incubated in the dark for 20-25 minutes at room temperature. Cells were washed and resuspended in PBS for analysis in the Beckman Coulter Cytoflex. Untreated cells were used to gate the debris, and false PI emission signals were gated out. To record the PI-positive cells, a B585/30 filter was used, and the percentage of cells of different stages were represented in a histogram. The percentages of cells of three biological repeats from each cell cycle stage are represented graphically.

### Quantitative RT PCR

Cells were cultured and harvested from confluent T25 flasks and the total RNA was isolated using Qiagen Total RNA isolation kit (Catalog No- 74104). Oligo-dT was used to synthesize cDNA by following the protocol of Promega Reverse transcription kit (Catalog No-A501). qPCR was performed using Powerup Sybr Master Mix (Applied Biosystem Catalog No.-A25742) for respective genes, primers are listed in **Table-S2**. Ct values were first normalized with the housekeeping control gene *GAPDH*. The fold change of the target genes expression against the empty vector treated control was calculated using the 2-ΔΔCt method and plotted graphically .

### m6A-RNA-Immunoprecipitation

m6A RNA Immunoprecipitation was performed using m6A antibody (Catalogue no.-E1611A). The beads were prepared with reaction buffer (150mM NaCl, 10mM Tris- pH 7.5 and 0.1% NP-40) and then incubated with antibody at 4°C for 1 hour. Followed by washing with reaction buffer using a Magnetic rack, beads-antibody complexes were incubated with 10µg of RNA at 4°C for one hour. Above solution was washed with a Reaction buffer, Low salt buffer (50mM NaCl, 10mM Tris- pH 7.5 and 0.1% NP-40) and High salt buffer (500mM NaCl, 10mM Tris- pH 7.5 and 0.1% NP-40) respectively using magnetic rack. Beads were resuspended with RLT buffer (Buffer of RNeasy Mini Kit, REF-74104) and eluted by using a magnetic rack. Eluted RNA was purified using nucleic acid binding beads (Zymo REF-A200-96). Followed by washing with 100% and 70% ethanol respectively, purified RNA was then eluted by NFW and then processed for cDNA synthesis and qPCR analysis. The target gene expression from the precipitated RNA was calculated by the percent input methods. The input (10%) was normalized to 100%, after which the fold change in expression was calculated for the treatment group (with antibody) relative to the control group (without antibody) for each cell line sample and subsequently represented graphically.

### Spheroid Growth Assay

Single cell suspension of MDA-MB- 468 cells was plated in 35mm culture dishes at a density of 1000 cells/ml under low attachment conditions in the spheroid media consisting of serum-free DMEM/F12 supplemented with 1% L-glutamine (Gibco), 1% penicillin/streptomycin (Gibco), N2 supplement (Gibco, 17502048,), B27 supplement (Gibco, 17504044), 10 ng/ml EGF (Invitrogen) and 20 ng/ml FGF (233-FB- 010/CF, R and D biosystems). The culture was grown for 5 days and supplemented with fresh medium. The spheroids were counted and documented on day 5 and day 7. Both the 5 days and 7 days spheroids were captured by the Zoe Fluorescent imager (Bio Rad), and analyzed using ImageJ software. The plate was divided into four quadrants and 7-10 images were captured from each part by selecting the most representative microscopic fields. The images were processed through ImageJ for measuring the sizes and counting the numbers. Both the day 5 and day 7 spheroid size and spheroid counts were plotted graphically.

### Spheroid Immunofluorescence assay

The spheroids were grown, counted and collected in a round bottom tube. The spheroids were then fixed with 4% Paraformaldehyde in suspension with constant agitation at 4°C for 15 min. The fixed spheroids were washed with 0.1% TBST and permeabilized with 0.5% TBST for 30 min at RT. The spheroids were then blocked in an antibody dilution buffer (2% BSA in 0.1% TBST) for 1hr at RT and subsequently incubated with primary antibody (1:250, Table-S1) overnight at 4℃. The spheroids were then washed for non-specific primary antibody binding and then incubated with Secondary antibody tagged with Alexa Fluor (1:500) for 3 hours at room temperature in the dark. The spheroids were washed, counterstained with DAPI, mounted and visualized under Zeiss fluorescence microscope. The images were processed and signals were quantified in ImageJ software. Total of 10-15 spheroids for each cell line were selected for fluorescence intensity calculation. ROI for each spheroid was selected with a background ROI and the fluorescence intensity was recorded using ImageJ software. Corrected fluorescence intensity was calculated using the mean fluorescence by normalizing the background intensity and then plotted in a graph.

### Gene expression analysis using “Genomic Data Commons” Data Portal (GDC)

In the GDC database, we built a cohort of breast cancer to study the gene expressions according to the requirements. The cohorts were built with following details- Programme-TCGA, Project-TCGA-BRCA, Primary site of cancer-Breast, Biospecimen-Tumor, Available dataset- RNA seq from Illumina platform, and molecular filter applied for TNBC (lack of estrogen, progesterone receptors and the lack of or low expression of the HER2), without molecular filter for all over Breast cancer. With the built cohort we proceeded to the “analyze center”. After studying the clinical analysis, mutation frequency analysis we proceeded to the gene expression clustering center. Gene input was provided according to our interest, which generates the heatmap representation, and the fold expression value for each sample was downloaded. We sort the value from higher to lower according to our gene of interest and then plot the set of gene expression from the same samples.

### Statistical Analysis

All experiments were performed with at least 3 biological replicates. Standard deviation and mean was calculated for the graph. Unpaired t-test was done to obtain the two-tailed p values, in GraphPad Prism. P values with significance levels are presented in the following way, P < 0.05=*, P < 0.01 =*\*\**, P <0.001 =***, and P <0.0001=****.

## Acknowledgments

The authors are grateful to the CAIF facility IISER Berhampur for imaging and flow cytometry facility, Sequencing Facility, ILS, Bhubaneswar.

## Author contributions

RSB designed the study, SD, SSP, RRB, SL, analyzed data, SD, RSB, SSP, RRB, SL, AMM, PD performed experiments, RSB, provided ideas and suggestions, RSB, supervised experiments, RSB, obtained funding, SD, RSB, SSP, SL wrote the manuscript with inputs from all authors.

## Abbreviations

m6A: N^6^ methyladenosine
TNBC: Triple negative breast cancer
WTAP: Wilms’ tumor 1 associated protein
FTO: Fat mass and obesity-associated protein
EMT: Epithelial to mesenchymal transition
CSC: Cancer stem cell
TME: Tumor microenvironment
KD: Knockdown

---

**Supplementary Figure 1:**
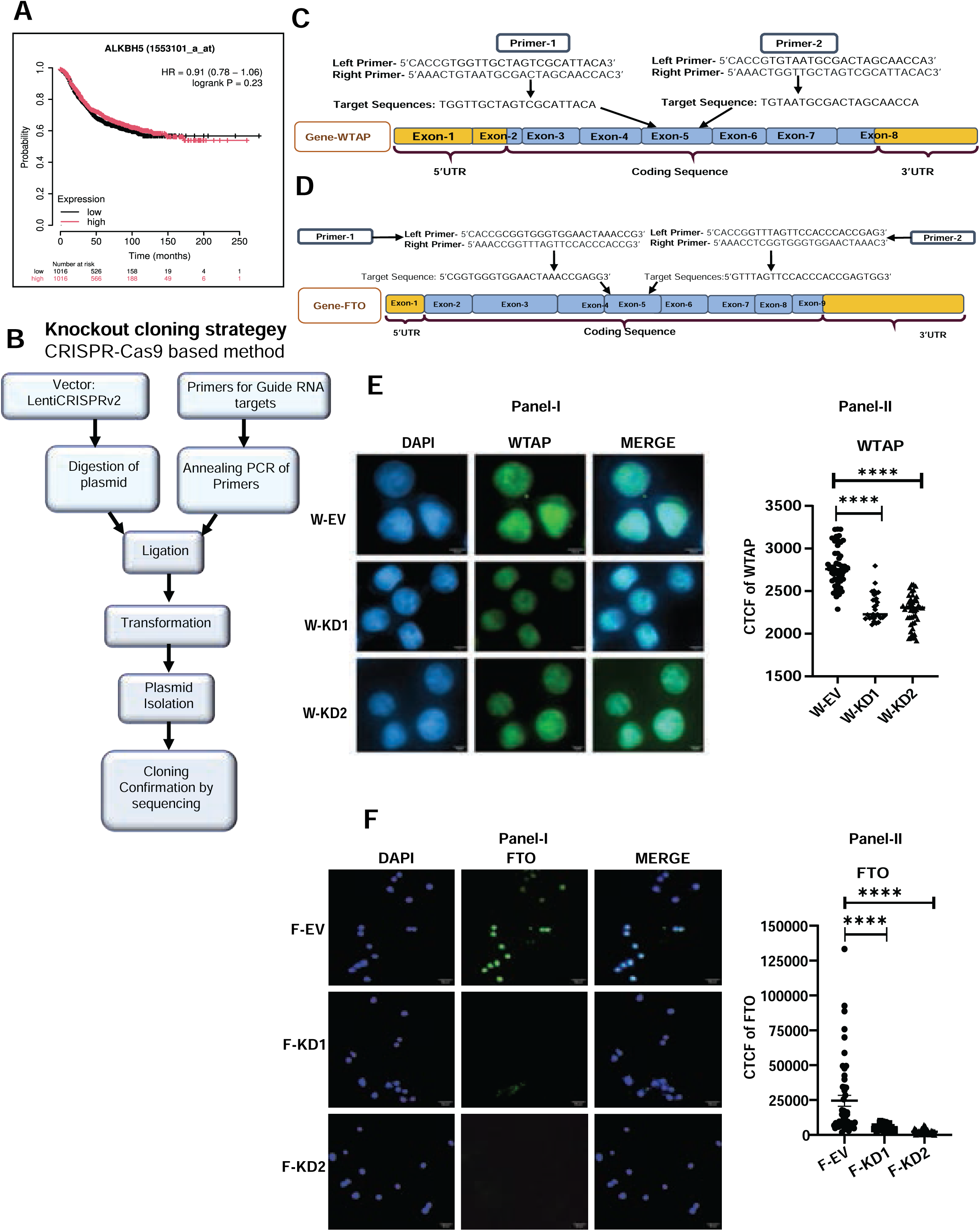
RNA Methylation modulators are differentially expressed in Triple Negative Breast Cancer Cells. **A.** Kaplan-Meier Plotter generated survival curves of breast cancer patients in ALKBH5 high and low expression conditions. **B.** Schematic representation of CRISPR-Cas9 based genome editing of m6A modifiers. **C**. The gene arrangement and guide RNA target details of WTAP on exon-5. **D**. The gene arrangement and guide RNA target details of FTO that target exon-5 of FTO. **E.** Immunostaining image (**Panel I**) of WTAP (Green Channel), DAPI (blue channel) and merged image of WTAP KD cells. Graphical representation of the corrected total fluorescence intensity calculated by ImageJ analysis (**Panel-II**). The error bars depict the SD and Mean, n=3, Significance analysis using student’s t-test performed by GraphPad, P-Value <0.0001=****. **F.** Immunostaining image (**Panel I**) of FTO (Green Channel), DAPI (blue channel) and merged image of FTO KD cells. Graphical representation of the corrected total fluorescence intensity calculated by ImageJ analysis (**Panel-II**). The error bars depict the Means and SD, n=3 Significance analysis student’s t-test performed by GraphPad, P-Value <0.0001=****.

**Supplementary Figure 2:**
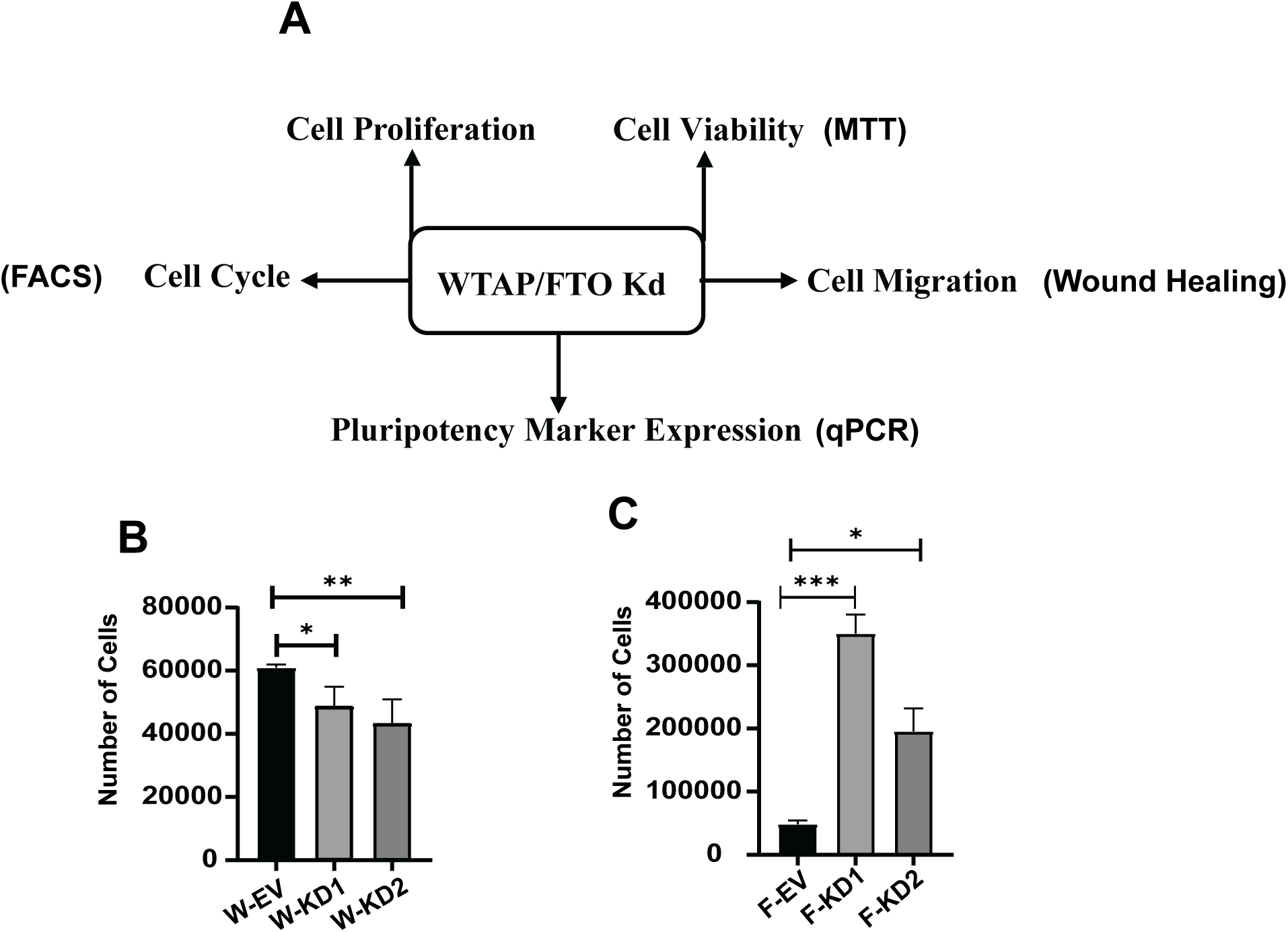
m6A Modifiers, WTAP and FTO Regulate Cell Proliferation in Triple Negative Breast Cancer Cells. **A.** Schematic representation of different assays to test biological functions regulated by WTAP/FTO. **B**. Graphical representation of cell proliferation in WTAP KD cells (W-KD1, W-KD2) with respect to empty vector (W-EV) control. The error bars depict the SD and Mean, n=3, Significance analysis using student’s t-test performed by GraphPad, P-Value < 0.05=*, < 0.01 =*\*\**. **C.** Graphical representation of cell proliferation in FTO KD cells (F-KD1, F-KD2) with respect to Empty vector control (F-EV). The error bars depict the Means and SD, n=3, Significance analysis student’s t-test performed by GraphPad, P-Value < 0.05=*, <0.001 =***.

**Supplementary Figure 3:**
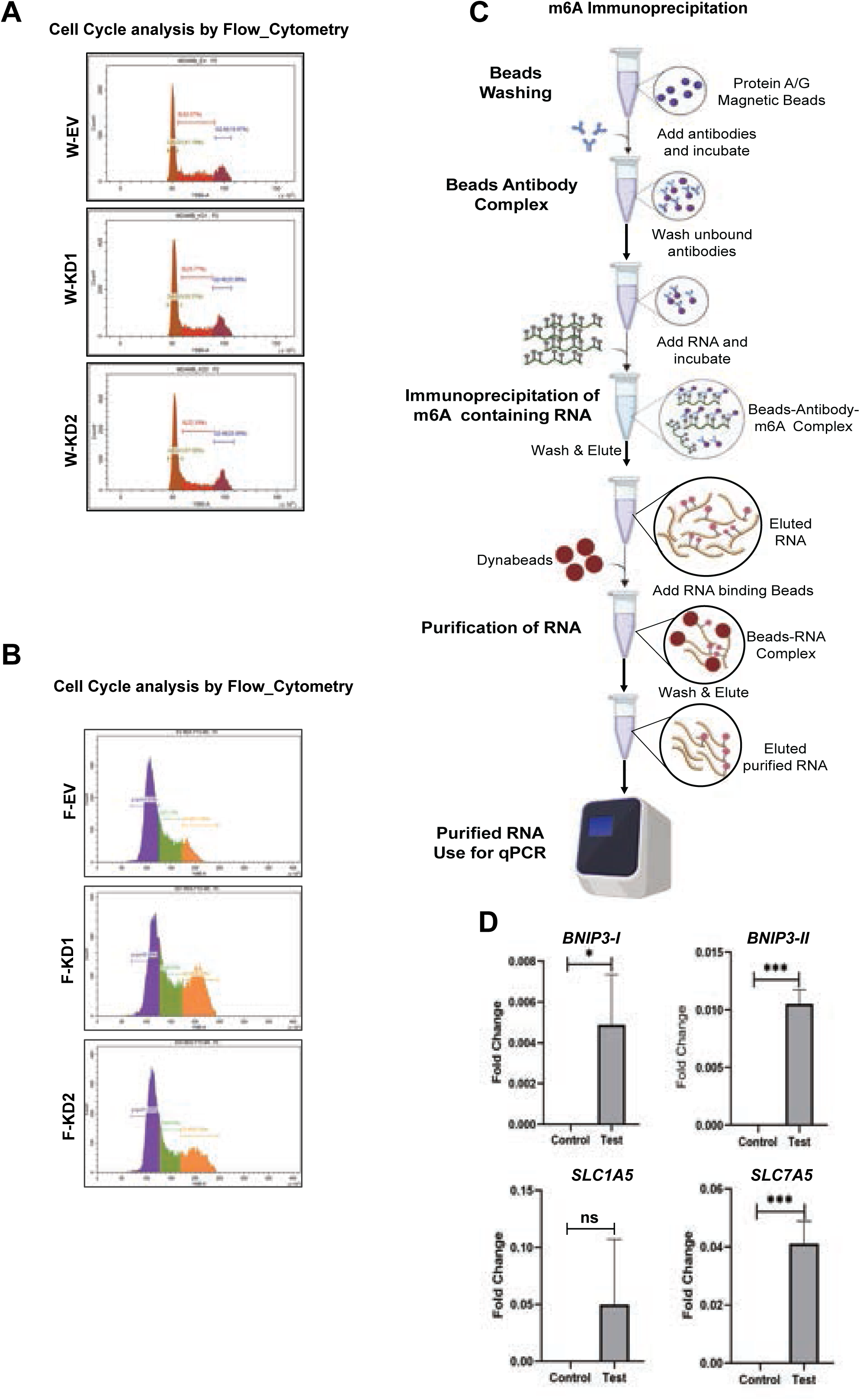
*Cyclin* expression is regulated in an m6A dependent manner in the TNBC cells. **A.** Histogram representation of flow cytometry-based analysis of cell cycle stages G0/G1, S, and G2/M for both knockdown and EV control in WTAP KD MDA-MB-468 cells **B.** Histogram representation of flow cytometry-based analysis of cell cycle stages G0/G1, S, and G2/M for both knockdown and EV control in FTO KD MDA-MB-468 cells. **C**. Schematic representation of the m6A-Immunoprecipitation analysis. **D**. Validation of m6A-IP by qPCR study of m6A enrichment on known FTO target genes (*BNIP3-I, BNIP3-II, SLC1A5, SLC7A5)*. The error bars depict the Means and SD, n=3, Significance analysis student’s t-test performed by GraphPad, P-Value < 0.05=*, <0.001 =***.

**Supplementary Figure 4:**
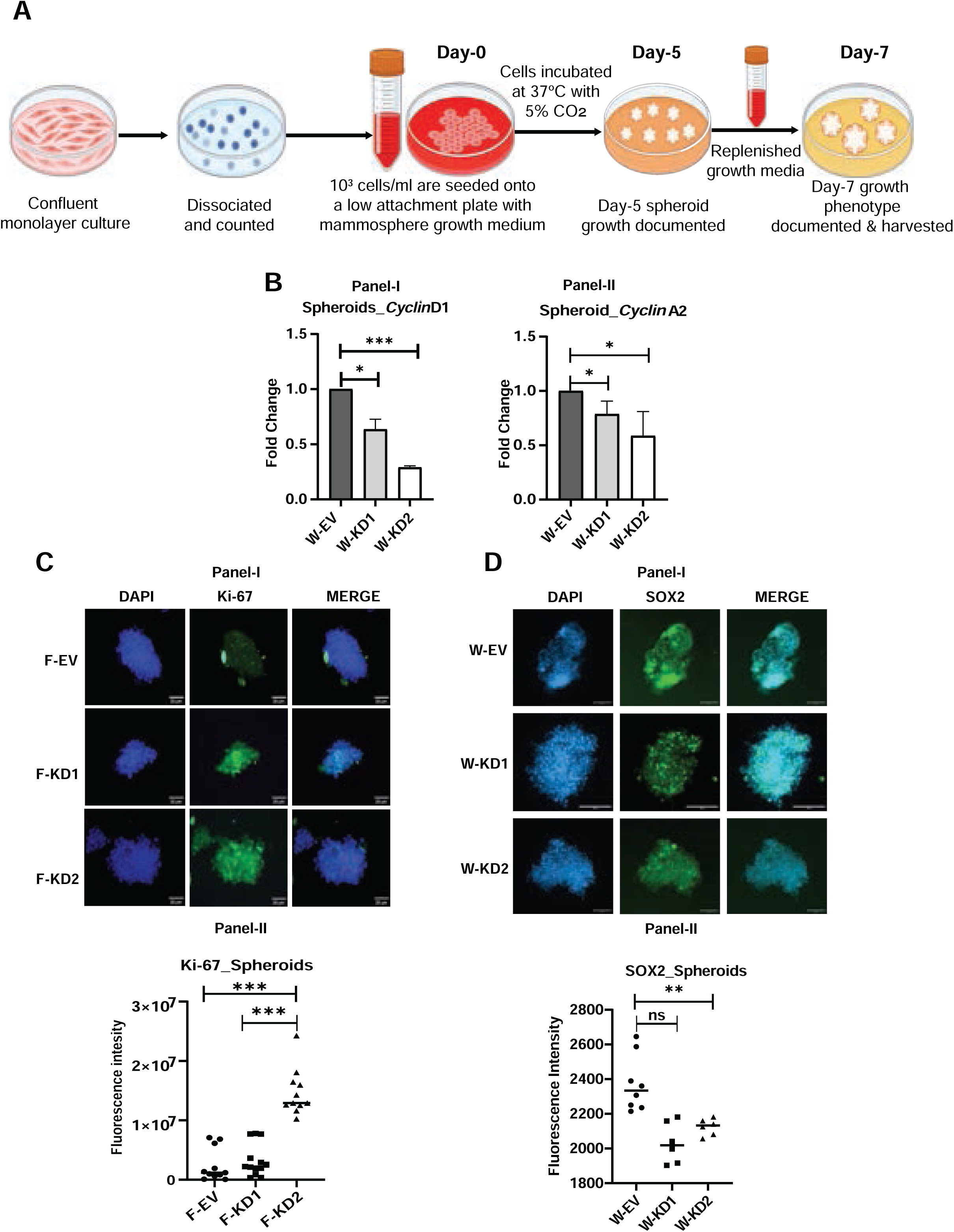
RNA Methylation modulates Cancer Stem Cell Maintenance in 3D spheroidal cultures of TNBC. **A**.Schematic of the spheroid culture procedure followed in this study. **B**. qPCR analysis of cell cycle regulator genes *Cyclin-D1* (Panel-I) and *Cyclin-A2* (Panel-II) in WTAP KD spheroid compared to EV control spheroid, normalised with housekeeping control *GAPDH*. The error bars depict the SD and Mean, n=3, Significance analysis using student’s t-test performed by GraphPad, P-Value < 0.05=*, <0.001 =***. **C**. The immunostaining image of Ki67 (Green Channel), DAPI (Blue Channel) and merged image of FTO KD spheroids (**Panel-I**). Graphical representation of the corrected total fluorescence intensity calculated by ImageJ analysis (**Panel-II**). The error bars depict the SD and Mean, n=3, Significance analysis using student’s t-test performed by GraphPad, P-Value <0.001 =***. **D** The immunostaining images of CSC marker SOX2 (Green Channel), DAPI (Blue channel), Merged image of WTAP KD spheroids. The mean fluorescence intensity was calculated by ImageJ analysis (**Panel-II**). The error bars depict the Means and SD, n=3, Significance analysis student’s t-test performed by GraphPad, P-Value < 0.01 =*\*\**.

**Supplementary Figure 5:**
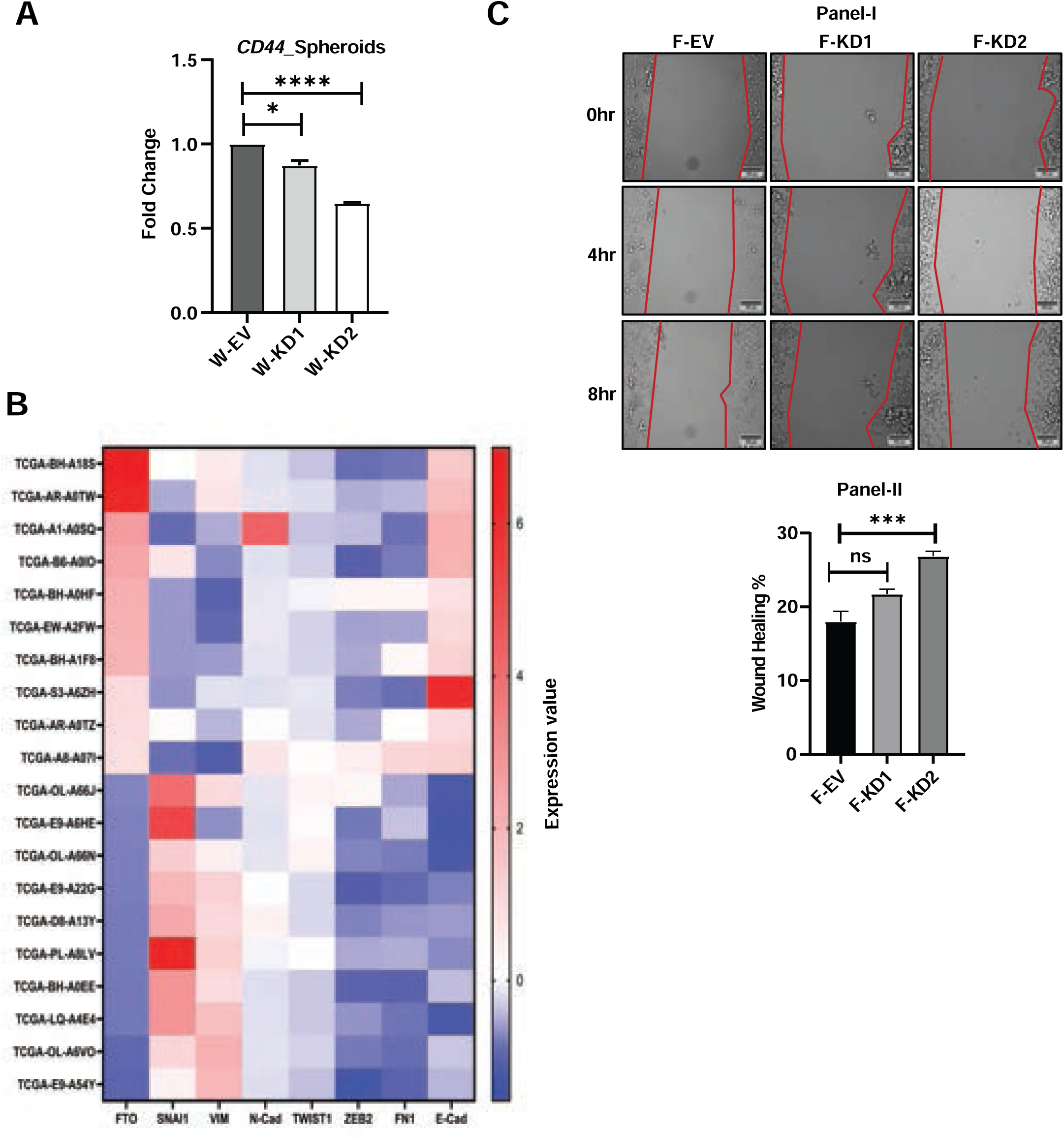
Epithelial Mesenchymal Transition associated markers and their alternative splicing contributes to tumorigenic potential of TNBC. **A.** qPCR analysis of stem cell marker genes *CD44* in WTAP KD spheroids compared to EV control, normalised with housekeeping control *GAPDH*. The error bars depict the SD and Mean, n=3, Significance analysis using student’s t-test performed by GraphPad, P-Value < 0.05=*, <0.0001=****. **B**. Heatmap of GDC data portal generated gene expression of EMT markers (*SNAI1, VIM, N-cadherin, TWIST1, ZEB2, FN1 and E-cadherin*) with respect to ascending order of FTO expression. Z-score transformed gene expression values are plotted. **C.** Brightfield image of wound healing assay at different time points (0 hours, 4 hours, and 8 hours) in FTO KD cells (F-KD1, F-KD2) compared to EV control (F-EV) (**Panel-I**). The percentage of cell migration was calculated by ImageJ and represented graphically (**Panel-II**). The error bars depict the Means and SD, Significance analysis performed by GraphPad, P-Value <0.001 =***.

**Supplementary Table-1:** Antibody Details.

| **Antibody** | **Catalog No.** |
| --- | --- |
| WTAP | ab155655 |
| FTO | ab127892 |
| Ki67 | ab15580 |
| SOX2 | ab137385 |
| KLF4 | CST #12173S |
| $\alpha/\beta$-Tubulin | CST #2148 |
| Anti-m6A | NEB E1611A |
| Alexa Fluor 488 | A32731 |
| Anti-rabbit IgG, HRP-linked<br>Antibody | CST 7074S |
| Anti-mouse IgG, HRP-linked<br>Antibody | CST 7076S |

**Supplementary Table-2:**
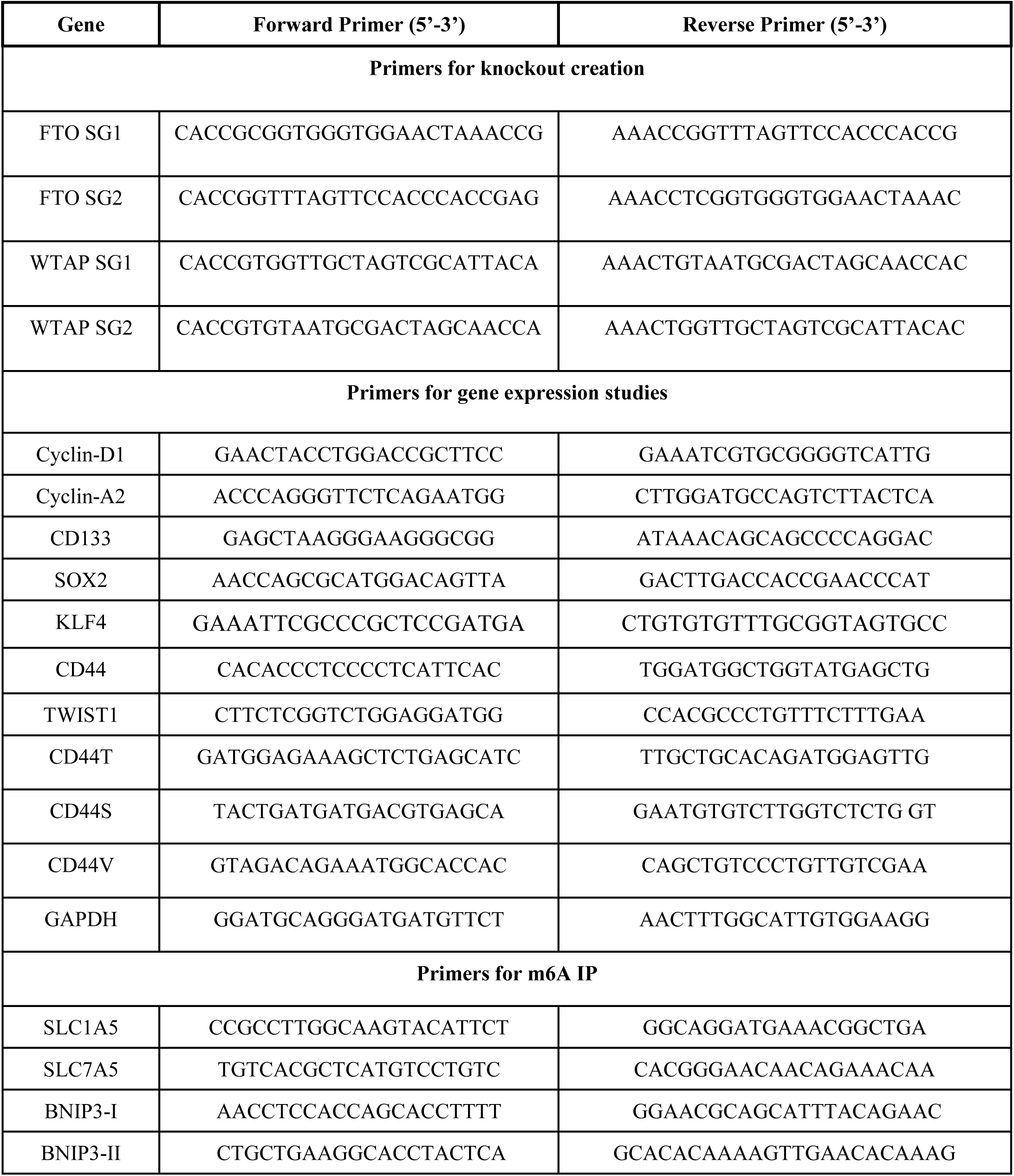
Primers Details.

